# Robust, sensitive, and quantitative single-cell proteomics based on ion mobility filtering

**DOI:** 10.1101/2021.01.30.428333

**Authors:** Jongmin Woo, Geremy C. Clair, Sarah M. Williams, Song Feng, Chia-Feng Tsai, Ronald J. Moore, William B. Chrisler, Richard D. Smith, Ryan T. Kelly, Ljiljana Pasa-Tolic, Charles Ansong, Ying Zhu

**Author notes:** Jongmin Woo and Geremy C. Clair contributed equally to this work. **Corresponding author:** Dr. Ying Zhu.

## Abstract

Unbiased single-cell proteomics (scProteomics) promises to advance our understanding of cell functions within complex biological systems. However, a major challenge for current methods is their ability to identify and provide accurate quantitative information for low abundance proteins. Herein, we describe an ion mobility-enhanced mass spectrometry acquisition and peptide identification method, TIFF (Transferring Identification based on FAIMS Filtering), designed to improve the sensitivity and accuracy of label-free scProteomics. TIFF significantly extends the ion accumulation times for peptide ions by filtering out singly charged background ions. The peptide identities are then assigned by a 3-dimensional MS1 feature matching approach (retention time, accurate mass, and FAIMS compensation voltage). The TIFF method enabled unbiased proteome analysis to a depth of >1,700 proteins in single HeLa cells with >1,100 proteins consistently quantified. As a demonstration, we applied the TIFF method to obtain temporal proteome profiles of >150 single murine macrophage cells during a lipopolysaccharide stimulation experiment and identified time-dependent proteome profiles.

## Introduction

Single-cell technologies have become the cornerstone of biomedical and cell biology research ^1, 2^. The emergence of single-cell RNA sequencing (scRNA-seq) and related single-cell sequencing technologies has illuminated unappreciated cellular heterogeneity and revealed cell subpopulations obscured in bulk measurements ^3^. However, many integrative studies have shown only low to moderate correlations between the abundance of RNA transcripts and their corresponding proteins ^4, 5^, as the translation of RNA into a functional protein can be affected by diverse events such as alternative splicing and microRNA regulation ^6^. Additionally, RNA measurements cannot infer post-translational modifications that modulate protein functions. Thus, there is an unmet need for broad proteome measurements at the single-cell level, which has lagged behind single-cell sequencing approaches.

Recent advances in sample preparation and mass spectrometry facilitate unbiased single-cell proteomics (scProteomics) ^7–20^. Microfluidic sample processing devices and systems have improved protein digestion efficiency and sample recovery by minimizing adsorptive losses ^13–16, 18, 20^. Tandem mass tag (TMT)-based isobaric labeling approaches (e.g., ScoPE-MS) have enabled multiplexed single-cell analysis in individual LC-MS runs ^7, 9, 11, 17, 19, 20^. The miniaturization of capillary electrophoresis or liquid chromatography has improved separation resolution and enhanced electrospray ionization efficiency ^21^. High-resolution MS analyzers combined with ion focusing devices, such as ion funnel, have increased detection sensitivity to the level where single molecules can be detected ^22^. State-of-the-art methodologies in scProteomics now can identify from ~700 to ~1,000 proteins from cultured single mammalian cells (e.g., HeLa) using label-free approaches ^8, 10, 14, 23, 24^ and from ~750 to ~1,500 proteins using TMT-labelling and signal boosting strategies ^7, 9, 11, 17–19^. Despite these advances, scProteomics remains immature, and significant technical challenges remain, including not only limited proteome depth and poor quantification performance, but also low system robustness for large-scale single-cell studies.

Because of the lack of a global amplification method for proteins, the coverage and quantification performance of scProteomics largely rely on the capabilities of MS measurement (e.g., sensitivity, speed, dynamic range). Although targeted MS measurements enable the detection of low copy number proteins and even single molecules ^22^, these measurements are generally performed using narrow m/z windows ^22^ or tandem mass spectra ^25^ to minimize background signals. Background ions, originating from ambient air and solvent/reagent impurities, dominate MS spectra during full m/z range acquisition. These abundant ions quickly fill ion trapping devices (e.g., ion trap or ion routing multipole) and limit the ability to trap ions over an extended time, which could otherwise accumulate more low-abundance ions of interest and improve detection sensitivity ^26, 27^. The high background signals generated by these ions can also significantly reduce the dynamic range of MS analyzers and deteriorate feature detection during downstream data analysis.

We reasoned that the removal of background ions should dramatically enhance the sensitivity of MS detection and improve the proteome coverage and quantitation performance of scProteomics. A variety of approaches have been developed to minimize background signals, including the use of a carbon filter in front of MS inlets to purify the ambient air ^28^, a picoliter-flow liquid chromatography (LC) system to reduce overall contaminates from air and solvent ^21^, a dynamic range enhancement applied to MS (DREAMS) data acquisition algorithm to reject highly abundant ions before ion accumulation ^27^, and a high field asymmetric waveform ion mobility spectrometry (FAIMS) interface to remove singly charged ions ^23^. Recently, Cong *et. al*. ^23^ demonstrated the coupling of FAIMS with low flow LC (20 nL/min) and Orbitrap Eclipse can identify ~1100 proteins from single cells. Because the peptides were identified by MS/MS, long LC gradients were required to collect sufficient numbers of MS/MS spectra for deep proteome coverages, which limited the analysis throughput. Herein, to address these challenges, we describe an MS1-centric data acquisition and peptide identification method, TIFF (Transferring Identification based on FAIMS Filtering), that significantly improves the proteome coverage, quantification accuracy, and throughput of label-free scProteomics. We demonstrated the capability and scalability of the TIFF method by studying macrophage activation with lipopolysaccharide (LPS) and by classifying dissociated human lung cells into distinct cellular populations.

## Results

### The TIFF method

The TIFF method is inspired by the accurate mass and time (AMT) tag approach ^29^, or other derivative approaches, such as “match between run” (MBR) implemented in MaxQuant ^30^ or IonQuant^31^, that generally rely on two measurements for the assignment of peptide identity: the accurate mass-to-charge ratio (m/z) and the LC retention time (RT). We have previously demonstrated that MBR improves the proteome coverage and reduces missing values in scProteomics ^13, 24^. The recent integration of ion mobility devices, including FAIMS at the interface between the LC system and mass spectrometer, provide an opportunity to use the additional ion-mobility separation dimension to reduce false discovery and improve coverage ^32^. We take advantage of this advance and utilize the FAIMS compensation voltage (CV) as a third matching feature (in addition to retention time and accurate mass) for peptide identification, as illustrated in Figure 1a. Briefly, a spectral library is constructed by repeatedly analyzing high-input samples on an LC-FAIMS-MS platform, with each LC-MS analysis utilizing a discrete FAIMS CV; in this case, CVs of −45V, −55V, −65V and −75V. Each peptide identified in the high-input analyses is associated with a unique 3-dimensional (3D) tag comprising LC retention time, accurate m/z, and FAIMS CV. Next, low-input samples (e.g., single cells) are analyzed by cycling through multiple FAIMS CVs (−45V, −55V, −65V, and −75V) within a single LC-MS analysis. A key aspect of the TIFF method is the mode of MS data acquisition, with most of the MS time spent on MS1 acquisition to enhance the accumulation of low-abundant peptide ions for sensitive detection. Compared with our previous FAIMS-based scProteomics method (Figure S1a and S1b),^31^ precursor ion sampling efficiency is increased by > 2 fold (Figure S1c). The fewer MS2 acquisitions generated within each cycle are sufficient to exploit the non-linear multi-sample alignment feature of MaxQuant. Subsequently, MS1 features in low-input samples (i.e., single-cells) are identified by matching to the spectral library and utilizing the unique 3D tag based on the MBR algorithm within MaxQuant ^30^.

**Figure 1.**
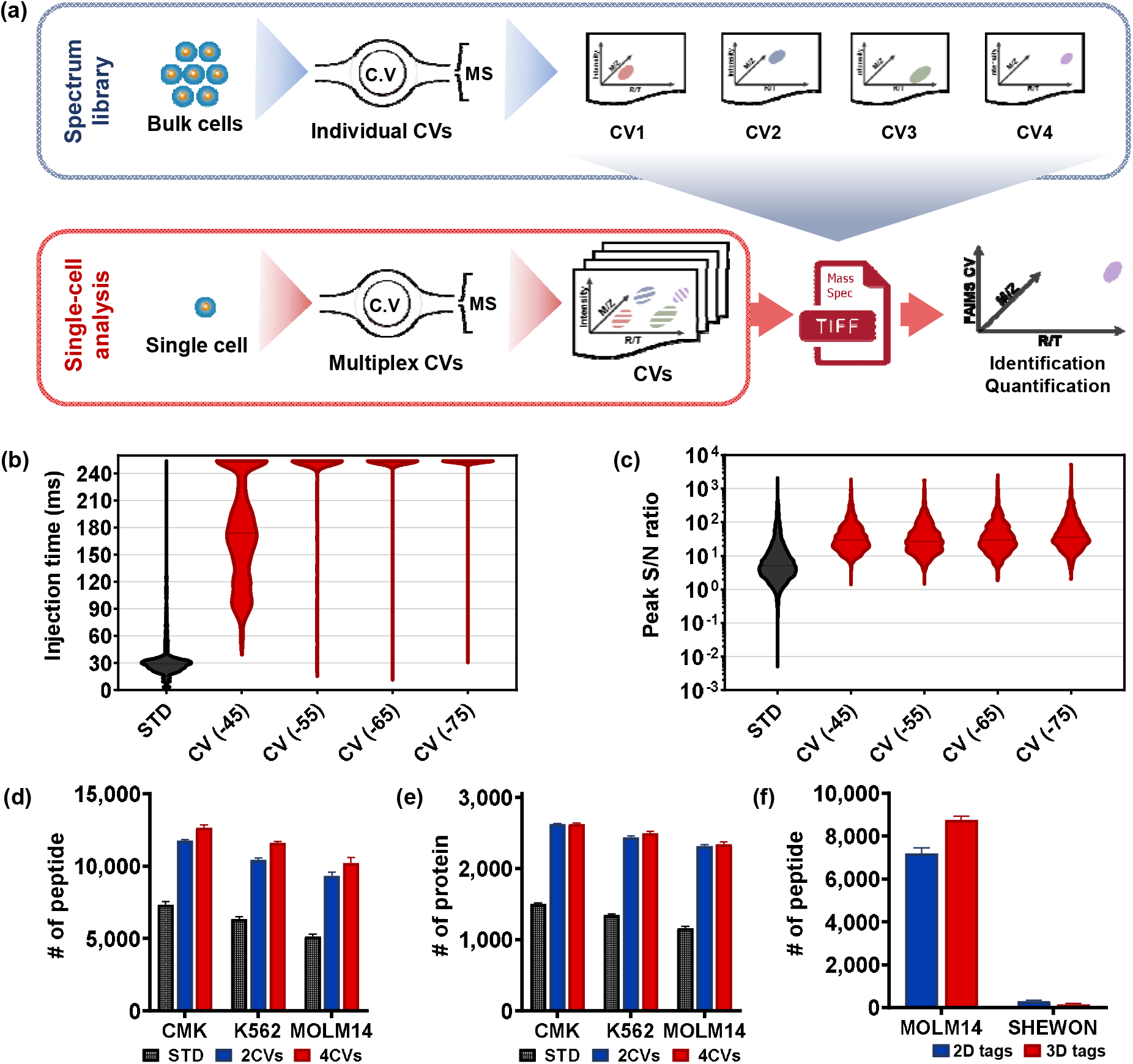
The concept of TIFF method. (a) Workflow of the TIFF method (Transferring Identification based on FAIMS Filtering). High-input samples (usually from 50 to100 cells) are analyzed by LC-FAIMS-MS with each LC-MS analysis utilizing a discrete FAIMS CV to generate a spectral library; Single-cell samples are analyzed by cycling through multiple FAIMS CVs for each LC-MS analysis. Peptide features in single cells are identified by matching to the spectral library based on three-dimensional (3D) tags, including retention time, m/z, and FAIMS CVs. (b) Injection time distributions of MS1 for the single-cell level peptides (0.2 ng, CMK cell) in the standard MS (STD, No FAIMS) method and FAIMS method with four different CVs. (c) The distributions of signal to noise ratios (S/N) of LC-MS features for the 0.2-ng peptides in STD run or FAIMS run with 4 CVs. (d-e) The average number of unique peptides and unique proteins using singlecell level (0.2 ng) peptide digests from three cell lines (CMK, K562, and MOLM14). Standards deviation error bars were obtained from the triplicate analysis. Benchmarking analysis was performed with the standard method, 2-CV TIFF (−45 and −65 V), and a 4-CV TIFF (−45, −55, −65 and −75 V) methods. (f) The number of human peptides (MOLM-14) and bacterial peptides (SHEWON) were identified from 2D and 3D tag methods. The bacterial peptides were considered as false identifications.

### TIFF improves LC-MS sensitivity

We first verified the utility of FAIMS to remove singly charged ions (“chemical background” noise) and create more “room” for peptide ion accumulation and enhance detection of low-abundance peptides. We analyzed single-cell equivalent amount (0.2 ng) of protein digests (CMK cells) with or without a FAIMSpro interface. Without FAIMS, most dominating signals correspond to singly charged ions, some of which are known to originate from plasticizers (e.g., m/z 391.28) and air impurities (e.g., m/z 445.12, 462.29, and 519.14) (Figure S2). Because these highly abundant contaminants quickly fill ion accumulation (or trapping) regions, the median ion injection/accumulation time is only 30 ms across the whole LC-MS analysis (Figure 1b). In comparison, when FAIMS is used, most dominating ion signals are multiply charged (Figure S2), and the median ion injection times increase from ~30 ms to ~180 ms for a CV of −45 V, reaching the maximal time of 254 ms for the other three CVs. This corresponds to an ~8.5× increase in ion sampling efficiency (Figure 1b). Benefiting from the low background and elongated ion accumulation, the median S/N of LC-MS features increased from 5.2 (STD) to 29.6 (FAIMS), representing a ~5-fold increase for all the CV values (Figure 1c).

To evaluate the improvements in MS sensitivity, we investigated several metrics related to proteome coverage, including the number of multiply-charged MS features, unique peptides, and proteins (Figure 1d-1e and Figure S3a-S3d). Briefly, we analyzed single-cell-level (0.2 ng) protein digests from three leukemia cell lines: CMK, K562, and MOLM14 with either a FAIMSpro interface or with a standard interface. Compared to the standard interface, the FAIMSpro interface and the TIFF method increased the number of multiply-charged MS features detected in the MS1 by > 3 folds (Figure S3a). Most of the increased peptide features appeared in the low-MS-intensity scale across all four FAIMS CVs (Figure S3e). Similarly, the TIFF method increased peptide identification by >75% (Figure 1d) and protein identification by >74% (Figure 1e). As expected, the MS/MS-based identifications were reduced due to the lower number of MS/MS scans (Figure S3b to S3d) in the TIFF method. Modulation of CVs within the TIFF method had a modest effect on the number of peptide features, peptides, and proteins, with only a slight increase using 4 CVs as opposed to 2 CVs. However, utilizing 4 CVs in the TIFF method yielded increases in summed peptide intensities compared to utilizing 2 CVs (Figure S3f), which subsequently improved the quantification performance as described below.

We evaluated whether the 3D feature matching approach could reduce false discovery rates by comparing it with the conventional 2D matching approach^29, 30^. We generated a mixed-species spectral library containing 20588 human peptides from MOLM14 cells and 9362 bacterial peptides from Shewanella Oneidensis MR-1. These Shewanella proteins were served as “decoy” proteins in the library. To do this, we analyzed 0.1-ng MOLM14 peptides with the 4CV-FAIMS method. During MaxQuant analysis with MBR algorithm, we either disabled or enabled the FAIMS CV matching function. As shown in Figure 1f and Figure S4, the conventional 2D matching approach results in total 7199 peptides identified, and 304 of them are bacterial peptides, representing a false matching rate of 4.1%. Encouragingly, when the 3D matching approach (TIFF) was applied, only 161 bacterial peptides were identified, corresponding to a false matching rate of 1.8 % (Figure S4). At the protein level, the false discovery rates of 2D and 3D matching approaches were estimated to be 10.8% and 5.3%, respectively.

### TIFF improves the quantification of scProteomics

Next, we evaluated whether the TIFF method improves quantification performance when compared with a standard approach. We compared the run-to-run reproducibility from triplicates using 0.2 ng of CMK cell digests with the standard, 2-CV TIFF, and 4-CV TIFF methods. While the distribution of the coefficients of variation was similar between the 2-CV TIFF and the standard methods, the median of the coefficients of variation for the 4-CV TIFF method was significantly reduced from 15.6% to 12% (Figure S5a). Such an improvement could be attributed to the enhanced sensitivity of the 4-CV TIFF method, allowing more low-abundance peptides to be identified. With the 4-CV TIFF method, > 80% of the proteins had no missing values and > 90% had no more than one missing value across the triplicates. Higher percentages of missing data were present with the 2-CV TIFF and standard methods (Figure S5b). To further assess the quantification accuracy of the 4-CV TIFF method, we performed a statistical analysis using samples from two cell types (CMK and K562). Proteins having at least 2 valid values in a given group were considered quantifiable. The 4-CV TIFF method exhibited a total of 2,345 quantifiable proteins that included ~98% of the proteins (1,052) using the standard method (Figure S5c). Because it was possible to quantify proteins more consistently with the TIFF method, we observed 1,053 differentially abundant proteins (DAPs) (FDR < 0.05 and S_0_=0.1) between the CMK and K562 cells, while only about half (i.e., 536 DAPs) were found using the standard method (Figure S6a-S6b). A total of 380 DAPs were shared between the two methods. As shown in Figure S5d, the linear correlation coefficient of protein fold-changes between the two label-free methods is high (R = 0.95). The slope of linear regression is ~1 (*K*), indicating similar fold changes between the two methods. Similarly, the 4-CV TIFF method showed improved quantification results over the standard method in the comparison between MOLM14 and the other two cell types (Figure S7 and S8).

### A streamlined label-free scProteomics platform

Having demonstrated that the TIFF method offers improvements in proteome coverage and quantification for mass-limited samples, we integrated it into our scProteomics pipeline that includes fluorescence-activated cell sorting (FACS) for cell isolation ^14^, a robotically addressed nanowell chip for single-cell processing (nanoPOTS, Nanodroplet Processing in One pot for Trace Samples) ^13^, a nanoliter-scale LC autosampler for reliable sample injection ^10^, and a low-flow liquid chromatography system (LC column with 50 μm i.d.) ^10^. Both single cells and pooled library cells can be isolated with FACS and processed with nanoPOTS. The integrated FACS-nanoPOTS-autosampler-TIFF-MS platform offers a complete solution from cell isolation to data acquisition and peptide identification for unbiased scProteomics, as well as other biological applications with mass-limited samples. The platform is robust and scalable. Since developed, it has been used to analyze > 1200 samples in our facility.

### Proteome coverage of single HeLa cells

We used HeLa cells to benchmark the TIFF-based scProteomics workflow. Using a tandem mass spectrometry approach (MS/MS), an average of 209 proteins were identified from single HeLa cells (Figure S9a). The number is comparable to our previously reported result (211 proteins) using a lower-flow LC-MS system (50 nL/min with 30 μm i.d. column) but without a FAIMS interface (Supplementary Table 1) ^14^, and 42% lower than that obtained using an ultra-low-flow LC system (20 μm i.d. column) and the newest generation (Eclipse) MS ^8^. The utilization of the 4-CV TIFF method dramatically increased the coverage to an average of 1,212 (± 10%) identified protein across 10 single cells (Figure S9a). The TIFF method doubled the total number of identifications compared with our previous report ^14^, reaching 1,771 unique proteins (Figure S9b). The number of identifications obtained with the TIFF method is comparable to the one we obtained using a 20-μm-i.d. column (20 nL/min)), a FAIMS interface, an Eclipse MS, and a long LC gradient ^23^.

The quantification consistency was also evaluated. Using protein iBAQ intensities, 684 out of 1,771 proteins had no missing values across the 10 HeLa cells (Figure S9c). 1,103 proteins were presented in at least 50% of the analyses. Pearson’s correlation coefficients had a median value of 0.95 between any two HeLa cells, indicating the high reproducibility of our integrated scProteomics pipeline (Figure S9d). Together, these results demonstrated that the integration of the TIFF method with high-efficiency singlecell preparation offers a sensitive and reliable scProteomics pipeline for label-free quantification.

### Preliminary application to dissociated primary cells from human lung

To initially explore the scProteomics platform for cell-type classification from dissociated primary cells, we analyzed non-depleted and non-labeled primary cells from the lung of a 2-year-old donor (Figure S10a). 19 single cells were processed and analyzed using the TIFF-based scProteomics workflow, resulting in a total of 986 identified proteins with an average of 390 identified proteins per single cell (Supplementary Table 2). We retained proteins identified in at least 8 of the 19 single cells (40% presence) for quantitative analysis, resulting in 402 quantifiable proteins (Supplementary Table 3). PCA analysis of the 402 proteins suggested the presence of at least three cell populations in the lung tissue single-cell suspension (Figure S10b).

To identify proteins distinguishing these populations, we performed the ANOVA test (permutation-based FDR < 0.05, S_0_ = 0), revealing 99 proteins (~20% of quantifiable proteins) that were differentially abundant across the three cell population groups/clusters (Supplementary Table 4) as visually represented in Figure S10c. Cell-type identity was assigned to each of the three cell population groups by comparing markers from the scProteomics data to lung cell type markers previously enumerated by bulk proteomics of sorted cell populations of human lung endothelial, epithelial, immune, and mesenchymal cells ^5^. Correspondence across the scProteomic and bulk proteomic markers revealed Cluster 1 represented a lung endothelial cell population, Cluster 2 represented a lung immune cell population, and Cluster 3 represented a lung epithelial cell population (Figure S11). For example, Caveolin-1 (CAV1) and Polymerase I and transcript release factors (PTRF), which were highly abundant in single-cell cluster 1 (Figure S10d and Figure S11), are known to structurally maintain the specialized lipid raft of caveola in lung endothelial cells ^33^. L-Plastin (LCP1) protein, important for alveolar macrophage development and antipneumococcic response ^34^, was highly abundant in bulk sorted immune cells as well as Cluster 2. Pulmonary surfactant-associated protein B (SFTPB), which facilitates alveolar stability by modulating surface tension ^35^ is known to be preferentially enriched in lung epithelial cells. SFTPB was highly abundant in bulk sorted epithelial cells as well as Cluster 3. The above results demonstrate the feasibility of the scProteomics platform for cell-type classification from non-depleted whole tissue single-cell suspension samples.

We also examined the abundance patterns of the 17 marker proteins based on scProteomics, bulk proteomics, and transcriptomics of the sorted populations (Figure S11). For the three protein markers mentioned above, we observed good agreement in all three measurement modalities. However, similar to our previous integrative study^5^, we also observed disagreement for some protein markers. For example, TUBB protein is identified as an endothelial cell marker in the proteomics dataset, but it is not significant in the transcriptomics dataset. In addition, among the 7 epithelial cell markers, only 1 protein/gene (SFTPM) is significantly expressed in both proteomics and transcriptomics datasets.

### Large-scale proteome profiling of single macrophage cells in response to lipopolysaccharide treatment

To further evaluate our platform for large-scale scProteomics analysis, we profiled proteome changes of single murine macrophage cells (RAW 264.7) after 24-hr and 48-hr lipopolysaccharide (LPS) stimulation relative to unstimulated cells (control) (Figure 2a). We analyzed a total of 155 individual RAW 264.7 cells, containing 54 unstimulated cells, 52 24-hr stimulated cells, and 49 48-hr stimulated cells. Our analysis identified a total of 1,671 proteins across the 155 individual cells. The median number of proteins identified per cell was 451. While lower than the number of proteins identified from single HeLa cells described above, we note that RAW 264.7 cells have a median diameter of 10 μm ^36^ compared to ~17 μm for HeLa cells ^12^; the 5-fold difference in cell volume likely accounts for the reduced coverages. We also observed control cells to have fewer identified proteins than LPS-stimulated cells. The median numbers of identified proteins were 307, 482, and 575 for control, 24-hr stimulation, and 48-hr stimulation, respectively (Figure 2b). Previous reports have indicated that stimulated RAW 264.7 macrophages increased in size and changed morphology upon LPS stimulation, potentially accounting in part for the difference in identifications ^36^. Of the 1,671 identified proteins, 519 were conservatively retained for quantitative analysis after filtering out proteins containing > 50% missing values in at least one experimental condition. Using a UMAP (the uniform manifold approximation and projection)-based dimensional reduction analysis^37^, the 155 individual cells partitioned into three distinct clusters on a two-dimensional plot correspond to the three experimental conditions (Figure 2c). Five stimulated cells (3 from 24 hrs and 2 from 48 hrs) are clustered into the control group, indicating a only small portion of RAW cells (~5%) are not sensitive to LPS stimulation.

**Figure 2.**
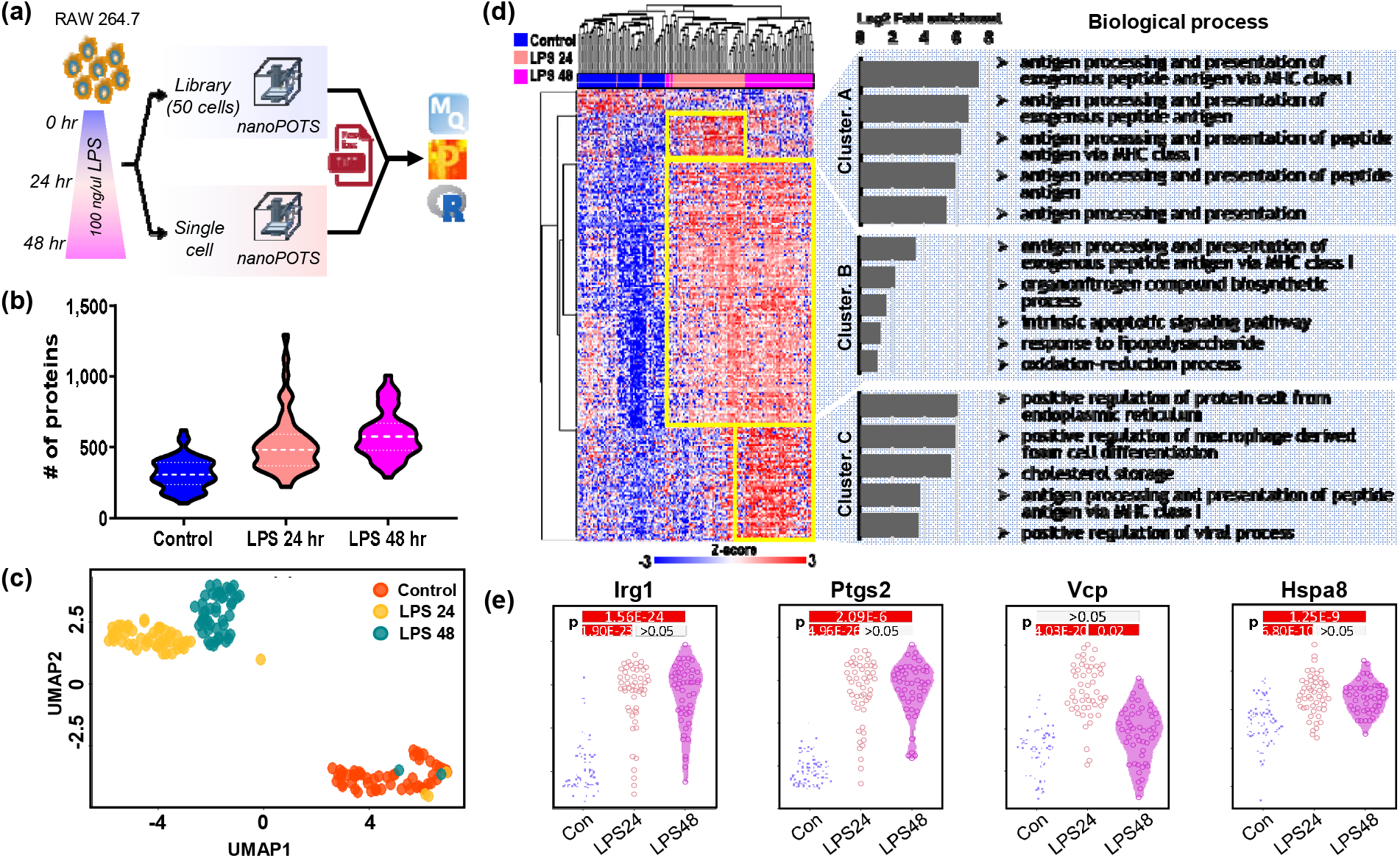
Single-cell proteomics analysis of time-dependent macrophage activation. **(a)** Illustration of workflow for scProteomics analysis of 155 macrophages containing untreated (control) cells and the cells treated by LPS for 24 and 48 hrs. **(b)** Violin plots of the distribution of the protein identification numbers for each treatment group. **(c)** UMAP projection showing the clustering of the 155 single macrophages cells based on treatment groups. **(d)** Heatmap showing the protein abundance differences across the 155 macrophage cells after statistical test using ANOVA (FDR <0.001, S0 = 5). The hierarchical clustering was performed using the Euclidean method for 250 DAPs by ANOVA test. Proteins in cluster A to C were applied to enrichment analysis using DAVID bioinformatics tools ^48^. **(e)** Abundance distributions of representative regulated proteins from different treatment conditions.

To identify the differentially abundant proteins (DAPs) that drive the partitioning of the three clusters, we performed an ANOVA test analysis (permutation-based FDR <0.001, S_0_=5). A total of 250 proteins were significantly modulated across the three groups (Figure 2d). Gene ontology analysis results showed that proteins increased in abundance at 24 hr LPS stimulation (Cluster A in Figure 2d) were primarily enriched in antigen processing and presentation processes (FE = 37.2 to 157.5, p < 0.01). Proteins increased at 24hr LPS stimulation and remained elevated through 48 hr LPS stimulation (Cluster B) were enriched in antigen processing and presentation (FE = 10.6, p < 0.05), response to LPS (FE = 2.5, p < 0.05), as well as oxidation-reduction (FE = 2.1, p < 0.01) processes, which are known to be a critical function of activated macrophage cells. Biological processes enriched in proteins increased after 48-hr LPS stimulation (Cluster C) included those related to protein exit from the endoplasmic reticulum (FE = 61.8, p < 0.05) and to foam cell differentiation (FE = 56.1, p < 0.05). The latter finding is in line with a previous report on the ability of LPS activated RAW 264.7 macrophages to differentiate into foam cells ^38^. Proteins associated with cholesterol storage (FE = 47.5, p < 0.05) were also increased in abundance after 48 hr LPS stimulation. Storage of cholesterol ester or triglyceride has been suggested to lead to the formation of foam cells ^39^.

Beyond functional enrichment analysis, our statistical analysis identified specific proteins previously described as being involved in the response process of macrophage cells to LPS stimulation. For example, immune responsive gene 1 (Irg1), known as a resistance-inducing protein against LPS ^40^, was up-regulated in macrophage cells exposed to LPS at both 24 and 48 hr (Figure 2e). Irg1 is highly expressed during various infections or TLR ligand stimulation in macrophages, which have been reported to regulate macrophage innate immune responses by controlling proinflammatory cytokines ^40^. Prostaglandin-endoperoxide synthase 2 (Ptgs2/Cox-2), an important precursor of prostacyclin enzyme which is expressed in macrophages exposed to LPS ^41^, was also significantly increased in LPS-stimulated macrophage cells (Figure 2e). Transitional endoplasmic reticulum ATPase (Vcp, also called p97) is involved in targeting and translocation of ubiquitinated proteins and the regulation of the inflammatory response in immune cells ^42^. We observed increased abundances of Vcp at 24 hr LPS stimulation with a decrease to basal levels at 48 hr LPS stimulation. The perturbation of cellular ubiquitin homeostasis supports the concept that variations in protein ubiquitination may be key to infection by pathogens and are also involved in triggering the defense mechanism of macrophages. Heat shock cognate 71 kDa protein (Hspa8), known to be involved in the presentation of antigenic peptides by major histocompatibility complex (MHC) class II (MHCII) molecules for CD4 + T cells, was significantly increased in LPS stimulated cells in line with previous studies that also showed this protein to be overexpressed in response to LPS stimulation ^43^.

## Discussion

In this study, we developed an ion mobility-enhanced MS acquisition and peptide identification method, TIFF (Transferring Identification based on the FAIMS Filtering), which was coupled with our previously described nanoPOTS scProteomics workflow ^10, 13^ to improve the sensitivity and accuracy of label-free scProteomics. MS acquisition efficiency was significantly improved by filtering out singly charged background ions and allowing ion accumulation for extended periods for sensitive detection. Compared with our previous FAIMS-based scProteomics workflow using an ultra low-flow LC column (20-μm-i.d.) and long gradient,^12^ the TIFF method dramatically improved both system robustness and analysis throughput to enable large-scale single-cell studies. The TIFF-based workflow enabled the identification of >1,700 proteins and quantification of ~1,100 proteins from single HeLa cells with label-free analysis. We demonstrated the robustness and scalability of the scProteomics workflow via a large-scale analysis of 155 single macrophage cells under different LPS stimulation conditions to reveal the biological processes at the single-cell level. Finally, we demonstrated the feasibility of classifying cell populations of a human lung.

While our label-free analysis of single cultured cells (e.g., HeLa) yielded >1000 proteins identified and similar numbers of proteins quantified, a similar analysis of single primary cells (e.g., human lung cells) resulted in the identification of significantly fewer proteins, presumably due to the fact that culture cells have larger sizes and more proteins mass. This again highlights the need to further improve the overall sensitivity of current scProteomics platforms to enable routine and deep single-cell proteome analyses of primary cells derived from tissues of animal models and human donors. One strategy for improving overall sensitivity is by further improving protein/peptide recovery. Sample recovery during sample processing procedures could be increased using smaller nanowells or low-binding surfaces to reduce adsorptive loss. Another strategy for improving overall sensitivity is through enhancing peptide separation resolution and ionization efficiency. With the advance of nanoLC pump technologies, the LC flow rates could be reduced to low nanoliter and to even picoliter-scale to further enhance peptide separation resolution and ionization efficiency. MS instrumentation with high ion-transmission optics and sensitive detectors could provide further enhancements in proteome coverage for single cells. In addition to FAIMS, other ion mobility-based technologies, including trapped ion mobility spectrometry (TIMS)^44, 45^, and particularly, structures for lossless ion manipulation (SLIM) can offer improved ion separation and overall ion utilization efficiencies. With all these developments, we believe the proteome depths of scProteomics will reach the level of single-cell RNA sequencing and ultimately become an indispensable tool in biological and medical researches.

## Acknowledgments

We thank the NIH-NHLBI Human Tissue Core (Dr. Gloria Pryhuber, Principal Investigator, U01 HL122700) for providing dissociated primary human lung cells and the family of the tissue donor for their generous and irreplaceable contribution to this research. We thank the insightful discussions from Aman Makaju at ThermoFisher Scientific (San Jose, CA) on the FAIMS interface. This work was supported by a Laboratory Directed Research and Development award (I3T) from Pacific Northwest National Laboratory (Y.Z.) and the NIH grants U01 HL148860 (C.A. and G.C.), R21 DC019753 (Y.Z.), and P41 GM103493 (R.D.S.). A portion of this research was performed on a project award (https://doi.org/10.46936/intm.proj.2020.51688/60000255) (Y.Z.) from the Environmental Molecular Sciences Laboratory, a DOE Office of Science User Facility sponsored by the Biological and Environmental Research program under Contract No. DE-AC05-76RL01830.

## Contributions

J. W., G. C., L. P. T., C. A., and Y. Z. proposed the method and designed the research. J.W., G. C., C.F.T., S. M.W, R. J. M., W. B. C, and Y. Z. performed cell culture, FACS sorting, proteomic sample preparation, and LC-MS experiments. J. W. and Y. Z. optimized the TIFF method. J.W., G. C., S. F., C. A., and Y. Z. analyzed the data. J. W., G. C., S. F., R. D. S., R. T. K., L. P. T., C. A., and Y. Z. wrote the manuscript.

## Data Availability

The mass spectrometry proteomics data have been deposited to the ProteomeXchange Consortium via the MassIVE partner repository with the dataset identifier MSV000085937.

## Materials and Methods

### Cell culture and single-cell sorting

All cell lines used in this study were maintained in a medium compatible with each cell line and incubated at 37 □ with 5% of CO_2_. Of the three Leukemia cell lines, K562 and MOLM14 cells were cultured in RPMI-1640 medium supplemented with 10% fetal bovine serum (FBS), and CMK cells were maintained in RPMI-1640 medium with 20% FBS added. For HeLa cells, DMEM supplemented with 10% FBS was added. RAW 264.7 cells were maintained in DMEM supplemented with 10% FBS followed to be stimulated with 100 ng/ul of LPS (Sigma Aldrich) in serum-free DMEM (Thermo Fisher Scientific) for 24 hr or 48 hr. For the control of RAW264.7 cells (non-treated), ten million cells were collected before stimulation with LPS. In the same way, LPS stimulated cells were harvested after 24 hr or 48 hr of treatments. HeLa and RAW 264.7 cells were washed by chilled PBS and sorted on the nanoPOTS chips (4 × 12, 1.2 mm diameter per well) using the Influx II cell sorter (BD Biosciences, San Jose, CA) as described previously ^14^. To build the in-depth spectral library, 50 cells of each cell line (or equivalent peptides of ~10 ng) were loaded onto the microPOTS chip (3 × 9, 2.2-mm diameter per well).

### Primary lung cells

The dissociated primary human lung cells was kindly provided by Dr. Gloria Pryhuber at University of Rochester Medical Center. The detailed protocol to generate the human lung cells was described previously^46^ and available on protocol.io (http://dx.doi.org/10.17504/protocols.io.biz5kf86). The dissociated lung cells in 90% FBS and 10% DMSO were cryo-frozen in −80°C freezer. A freezing vial was shipped to PNNL on dry ice. The cells were thawed and resuspended in DMEM with 10%FBS for 1 Hr prior to be centrifuged at 800 g for 10 min. The supernatant was removed and cells were washed in DPBS. To gate out dead cells or cell debris, the cells with labeled with Calcein AM viability dye (Thermo Fisher). Similar to the FACS-sorting procedures above, we sort 50 cells into microPOTS chips for library generation and single cells into nanoPOTS chips for analysis.

### Protein digestion

For the low-input mock samples (0.2 ng, equivalent amount peptides to a single-cell), AML cell lines were lysed in a tube with lysis buffer including 50 mM NH_4_HCO_3_ (pH8.0), 8 M UREA, and 1 % phosphatase inhibitor followed by sonicated in a cold bath for 3 min. After the measurements of the protein concentrations by BCA assay (Thermo Fisher Scientific), proteins equivalent to 200 μg were reduced in 5 mM DTT for 1 hr at 37 □ and alkylated with 10 mM iodoacetamide (IAA) in the dark for 1 hr at room temperature. Eight-fold diluted samples with 50 mM NH_4_HCO_3_ were digested with Lys-C peptidase at 37 □ with a ratio of 50:1 (w/w) for 3 hr followed by digesting with trypsin with a ratio of 50:1 (w/w) at 37 □ overnight. The tryptic digested peptides were acidified by 0.5% trifluoroacetic acid (TFA) at final concentration, then desalted using C18 SPE tips. After concentrated, the BCA assay was performed to estimate the final concentration of the peptides. Using the nanoPOTS robot, 0.2 ng and 10 ng of the peptides from each AML cell line were loaded on the nanowell/microwell chips and completely dried by a vacuum system ^10^.

For single-cell analysis, single and 50 FACS-sorted cells on the chip were processed on the nanoPOTS platform for single cells and spectral library, respectively. To extract proteins, we first added a lysis buffer containing 0.2% n-Dodecyl b-D-maltoside (DDM) and 5 mM DTT in 0.5× PBS and 25 mM NH_4_HCO_3_ buffer in each well, then incubated for 1 hr at 70 □. Denatured and reduced proteins were alkylated with 10 mM IAA in the dark for 30 min at RT. Double enzymatic digestions were performed by incubating with LysC (1 ng for single-cell, 5 ng for 50 cells) for 4 hr at 37 □ followed by treatment with trypsin (2 ng for single-cell, 10 ng for 50 cells) overnight. Peptides were acidified with 5% formic acid and completely dried using a vacuum system. All chips were stored in a −20 □ freezer until MS analysis.

#### Shewanella oneidensis

MR-1 peptide was obtained from a non-related study. The sample preparation procedures were described in detail previously^21, 47^.

### LC-FAIMS-MS/MS analysis

In-house assembled nanoPOTS autosampler with an in-house packed SPE column (100 μm i.d., 4 cm, 5 μm, 300 Å C18 material, Phenomenex) and an LC column (50 μm i.d., 25 cm, 1.7 μm, 190 Å C18 material, Waters) heated to 50 □ using AgileSleeve column heater (Analytical Sales and services, Inc., Flanders, NJ) was used for sample analysis ^10^. Briefly, samples were dissolved with Buffer A (0.1% formic acid in water) on the chip, then trapped on the SPE column for 5 min. After washing the peptides, samples were eluted at 100 nL/min and separated using a 60-min gradient from 8% to 35% Buffer B (0.1% formic acid in acetonitrile).

An Orbitrap Fusion Lumos Tribrid MS (Thermo Scientific) operated in data-dependent acquisition mode was used for all analyses. Peptides were ionized by applying a voltage of 2,000 V or 2,400 V for standard or FAIMS methods, respectively.

For the standard method, precursor ions with mass range 375-1600 m/z were scanned at 120,000 resolution with an ion injection time (IT) of 254 ms and an AGC target of 1E6. To analyze pooled samples for generating the spectral libraries, the selected precursor ions with +2 to +7 charges were fragmented by a 30% level of high energy dissociation (HCD) and scanned at 60,000 resolution with an IT of 118 ms and an AGC target of 1E5. When single-cell level (0.2 ng) peptides were injected, fragmented peptide ions were scanned at 120,000 resolution with an IT of 246 ms and an AGC target of 1E5.

For the TIFF method, the ionized peptides were fractionated by the FAIMSpro interface using a 2-CV (−45, −65 V) method or a 4-CV (−45, −55, −65, −75 V) method. Fractionated ions with a mass range 350-1500 m/z were scanned at 120,000 resolution with an IT of 254 ms and an AGC target of 1E6. For the pooled samples for generating a spectral library, a single CV was used for each LC-MS run. Precursor ions with intensities > 1E4 were selected for fragmentation by 30% HCD and scanned in an Ion trap with an AGC of 2E4 and an IT of 150 ms. For single-cell samples, cycle times of 1.5 s and 0.6 s were used for the 2-CV and 4-CV methods, respectively. Precursor ions with intensities > 1E4 were fragmented by 30% HCD and scanned with an AGC of 2E4 and an IT of 254 ms.

### Data analysis

All raw files were processed by MaxQuant (Ver. 1.6.2.10) with the Uniport protein sequence database of *homo sapiens* (Downloaded in 03/12/2020 containing 20,364 reviewed sequences) and of *mus musculus* (Downloaded in 5/19/2020 containing 17,037 reviewed sequences) using the Andromeda search engine with a 6-ppm precursor ion tolerance after mass calibration ^30^. Protein acetylation in N-terminal and oxidation at methionine were chosen as variable modifications. Carbamidomethylation of cysteine residues was set as a fixed modification. Both proteins and peptides were filtered with a false discovery rate (FDR) less than 0.01. Match between runs algorithm in Maxquant was activated with a matching window of 0.4 min and alignment windows of 10 min. For raw files with multiplex FAIMS CVs, we converted them to multiple mzxml files corresponding to separate individual CVs using an in-house converting tool (https://github.com/PNNL-Comp-Mass-Spec/FAIMS-MzXML-Generator/releases). Those separated files were assigned to non-adjacent fractionation numbers (e.g., 1, 3, 5, 7) during the Maxquant search to ensure feature matching only occurs between the files with the same CV.

For label-free quantification of single-cell-level peptides (0.2 ng) for three AML cell lines and dissociated human lung single-cell, Perseus (Ver. 1.6.12.0) was utilized for the data clean and statistical analysis. The iBAQ algorithm was used for the single-cell analysis because the iBAQ values are proportional to the molar quantities of the proteins. We log2 transformed the iBAQ values after filtering out contaminants and reverse identifications. Missing values were imputed based on a standard distribution (width: 0.3, downshift: 1.8) to simulate signals for low-abundance proteins. Data were normalized using width adjustment, which subtracts medians and scales for all values in a sample to show equal interquartile ranges. Two-way t-tests were performed for the pairwise comparison of the AML cell lines proteomes utilizing the threshold of Benjamini-Hochberg FDR < 0.05 and S_0_=0.1, while ANOVA tests were employed for multiple sample tests of dissociated human lung single cells with Permutation based FDR < 0.05. To clarify cell populations from dissociated lung cells, multiple steps including principal components analysis (PCA) and hierarchical clustering were employed using Perseus. Gene ontology analysis for the biological process of the molecules was performed in DAVID web-based bioinformatic tools (database version 6.8, https://david.ncifcrf.gov/summary.jsp).

The processing of the macrophage single-cell data was performed using an R package; RomicsProcessor v1.1.0 (https://github.com/PNNL-Comp-Mass-Spec/RomicsPro). Briefly, the “proteingroups.txt” output file of the MaxQuant search was imported as a multilayered R object with its associated metadata to extract iBAQ values of the identified proteins. The iBAQ values were then log2 transformed and filtered to allow maximal missingness of 50% within at least one given condition. After median normalization, batch correction was applied to remove the batch effects between chips using ComBat algorithm from the SVA package (v3.36.0). The missing values were imputed using the function of imputeMissing() and UMAP (the uniform manifold approximation and projection)-based dimensional reduction analysis was performed using the romicsUmapPlot() function in the RomicsProcessor package. For the statistics, ANOVA test was applied with a Benjamini-Hochberg FDR < 0.001 and a S_0_=5; we applied a highly significant level to a large number of macrophage cells data in which the group was clearly distinguished by the duration of LPS treatment to give a statistical role to the difference between the median value.

## Supplementary Materials

Figure S1. Comparison of different MS acquisition methods.

Figure S2. Representative MS raw spectra obtained with and without FAIMS interface.

Figure S3. Benchmarking of the detection sensitivity using different MS acquisition methods.

Figure S4. Evaluation of false matching rates by matching a human sample to a mixed-species spectral library containing both human and bacterial peptides.

Figure S5.The evaluation of the quantification performance of the TIFF method.

Figure S6. Differentially abundant proteins between CMK and K562 cell lines obtained from standard and TIFF methods.

Figure S7. Differentially abundant proteins between K562 and MOLM14 cell lines.

Figure S8. Differentially abundant proteins between CMK and MOLM14 cell lines.

Figure S9. ScProteomics of HeLa cells using TIFF method.

Figure S10. ScProteomics for classifying cell populations of a human lung.

Figure S11. Comparison of quantitative protein markers for human lung cells.

Supplementary Table 1. Numbers of identified proteins in single mammalian cells from previously published papers using nanoPOTS and label-free analysis.

Supplementary Table 2. List of identified proteins from 19 single lung cells.

Supplementary Table 3. List of 402 quantifiable proteins of 19 single lung cells.

Supplementary Table 4. A list of statistically significantly abundant proteins classifying three cell populations.

