## Supplementary_Information for "Robust, sensitive, and quantitative single-cell proteomics based on ion mobility filtering"

Woo et al.

**Supplementary Materials**

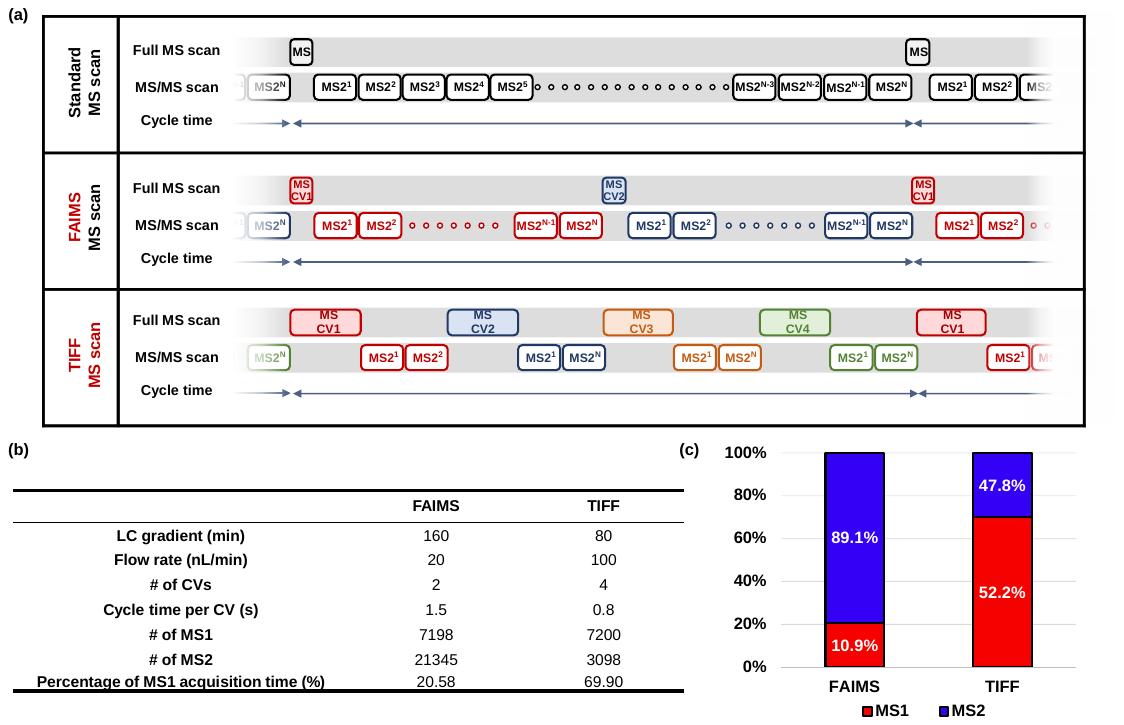

**Figure S1.** **(a)** Schematic illustration of three different MS acquisition methods. (Upper) standard MS method without FAIMS; (Middle) standard FAIMS-MS method; (Bottom) The transferring identification based on FAIMS filtering (TIFF) method. In the TIFF method, the elongated ion accumulations for MS1 scan can increase the sensitivity of MS1-level peptide detection. The peptide features are identified by matching to a spectral library based on 3D tags (LC retention time, accurate m/z, and FAIMS CV). Small number of MS/MS scans are used for non-linear alignment during MaxQuant search. **(b)** The comparison of standard FAIMS^1^ with the TIFF method for single-cell proteomics. **(c)** The normalized MS acquisition time for MS1 and MS2 scans. The precursor ion sampling efficiency of the TIFF method is increased by > 2 folds compared with the previous FAIMS method^1^.

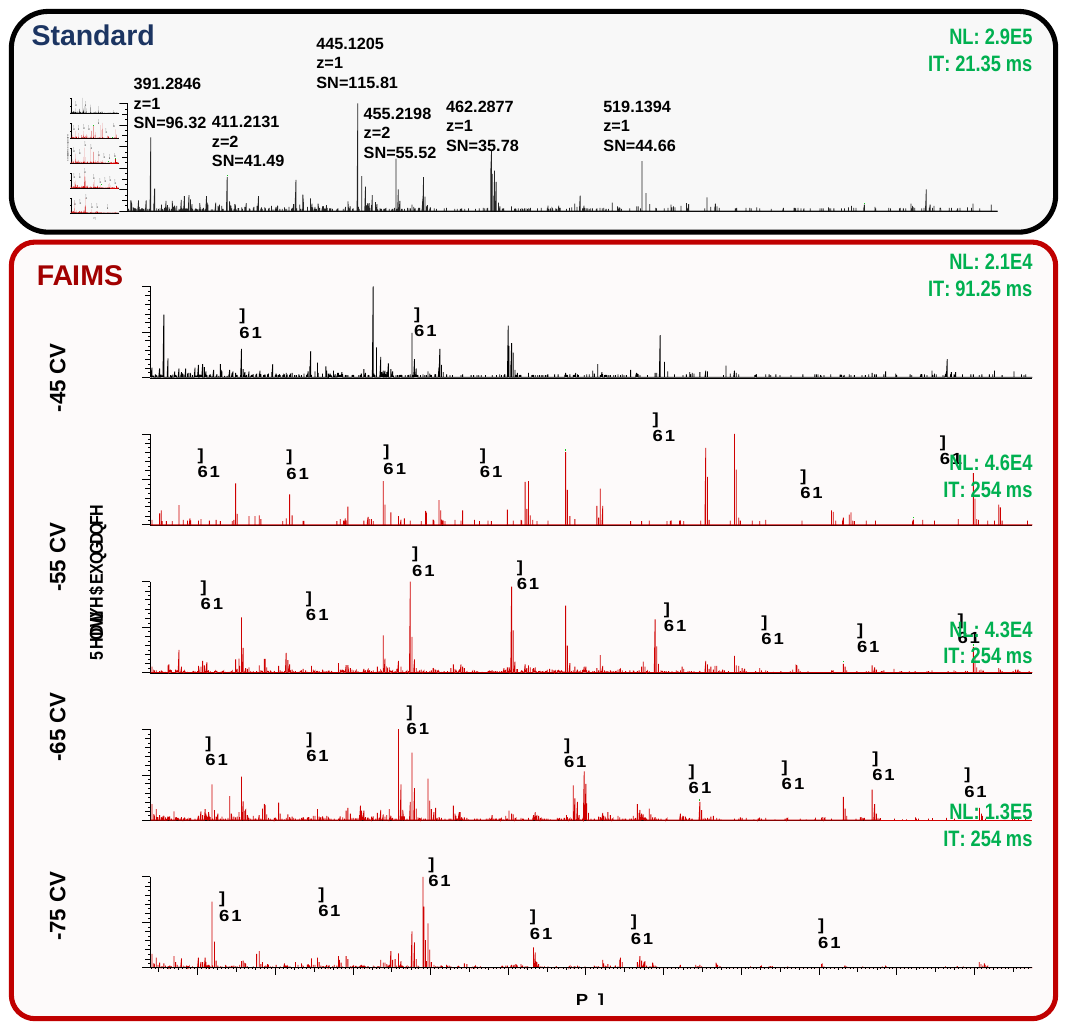

**Figure S2.** Representative spectra are chosen from the MS raw files of a standard method (in the blue box) and a FAIMS method with 4 CVs (in the red box). The spectra are extracted from a similar retention time. Spectra are labeled with m/z, ion charge state, and signal to noise (SN) value.

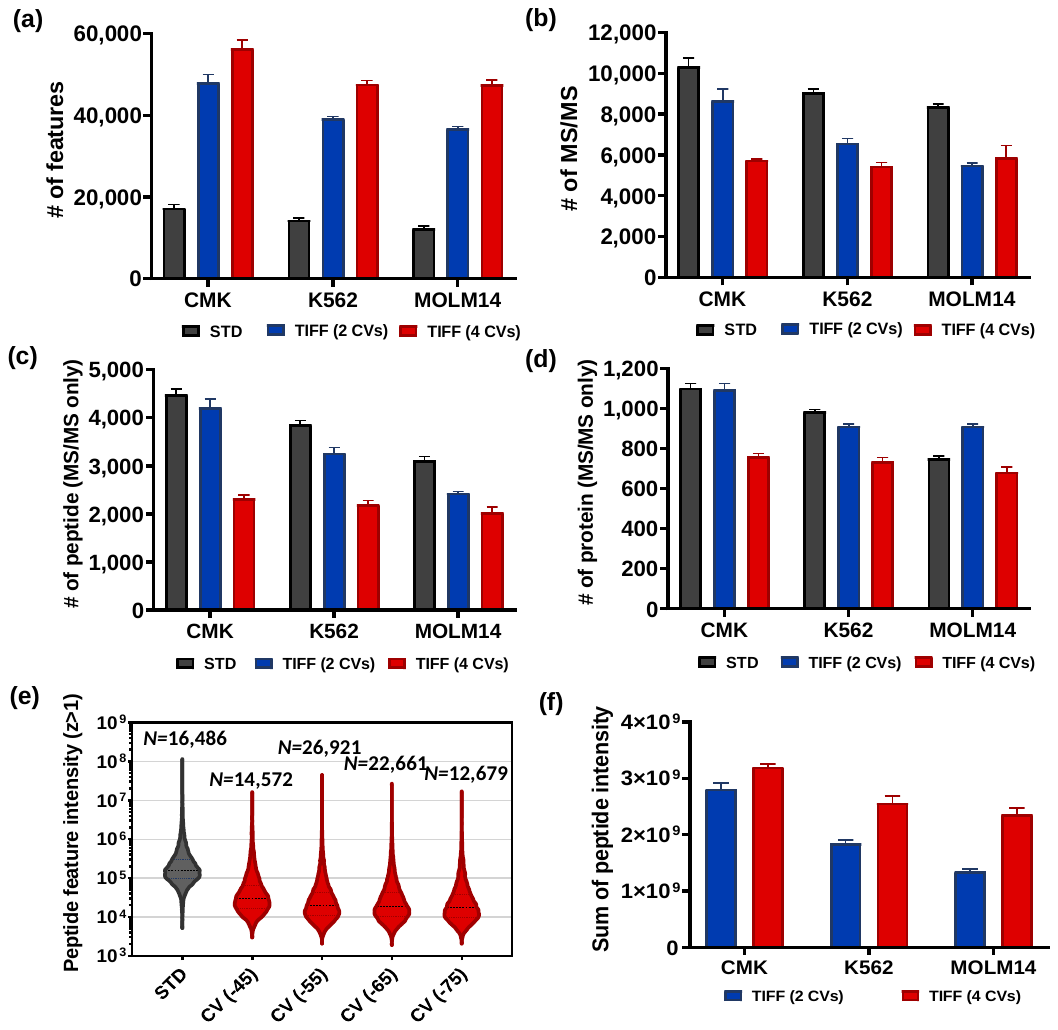

**Figure S3.** **(a-d)** Benchmarking of the standard, 2-CV-TIFF, and 4-CV TIFF methods using single-cell level peptides (0.2 ng) from three cell lines (CMK, K562, and MOLN14). **(a)** The number of peptide features (charge>+1); **(b)** MS/MS scans; and **(c)** unique peptides; **(d)** proteins identified by MS/MS. **(e)** Intensity distributions of peptide features (z > +1) obtained by the standard and 4-CV-TIFF methods using 0.2-ng CMK peptides. Labeled numbers indicate the numbers of detected peptide features. An in-house MASIC tool was used to select the peptide features from MSGF+ results. **(f)** The summed peptide intensities from the 2-CV and 4-CV TIFF methods. Error bars indicate standard deviations obtained from the triplicate analysis.

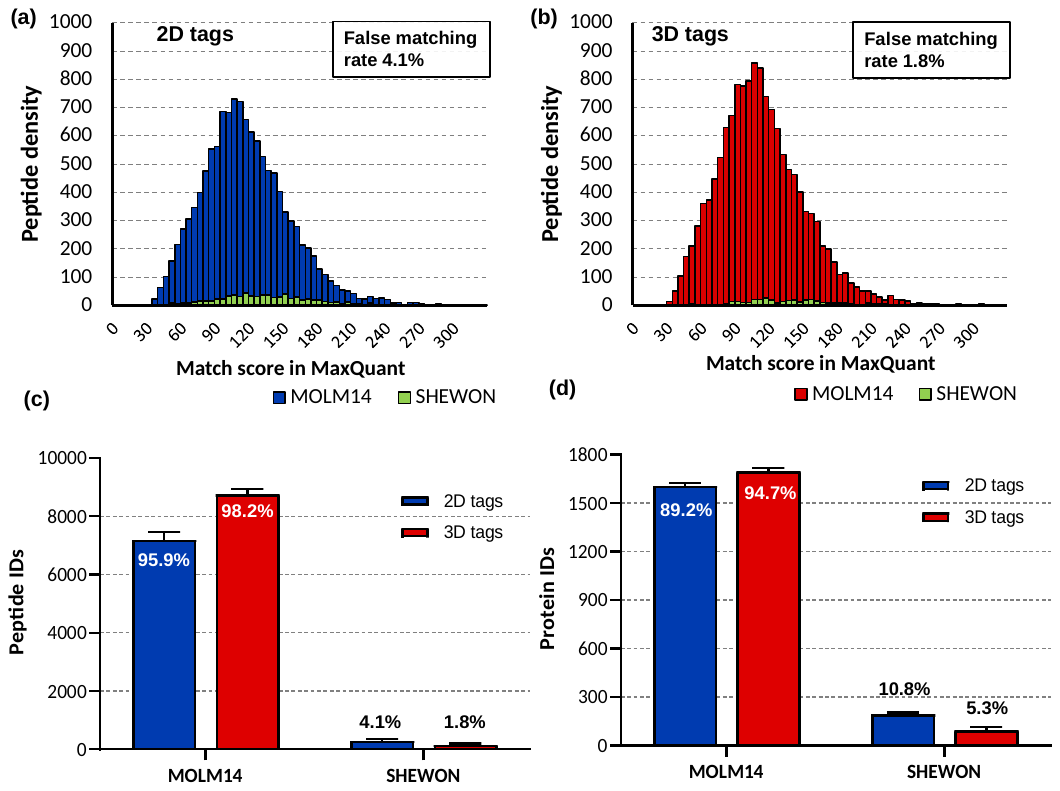

**Figure S4.** Evaluation of false matching rates by matching a human sample to a mixed-species spectral library containing 20588 human peptides from MOLM14 cells and 9362 bacterial peptides from Shewanella Oneidensis MR-1. Histogram of the number of identified peptides with **(a)** two-dimensional tags (m/z and RT), or **(b)** three-dimensional tags, (m/z, RT, and FAIMS CV). **(c)** False discovery rates at the peptide and **(d)** protein levels using 2D or 3D matching approaches.

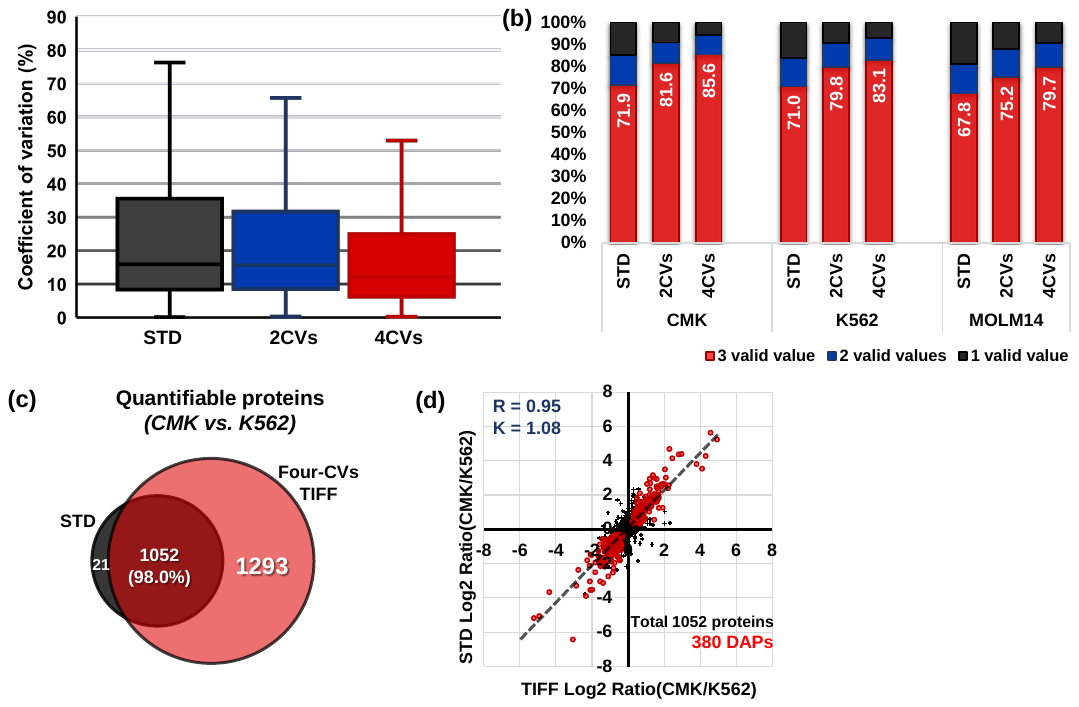

**Figure S5.** **The evaluation of quantification performance of the TIFF method. (a)** Distributions of the coefficient of variations for quantified proteins using the three MS acquisition methods (STD, 2-CV-TIFF, and 4-CV-TIFF). **(b)** Percentage of valid values using the three methods. Red, blue, and black colors indicate the percentages of proteins with valid values of 3, 2, and 1 across the triplicate, respectively. **(c)** Overlap of quantifiable proteins between CMK and K562 samples measured by standard and 4-CV-TIFF methods. **(d)** The linear correlation of log2-transformed fold changes of CMK and K562 cells between the TIFF method (4 CVs) and the STD method. Red dots indicate differentially abundant proteins (DAPs) in both methods calculated by t-test (FDR<0.05, S_0_=0.1).

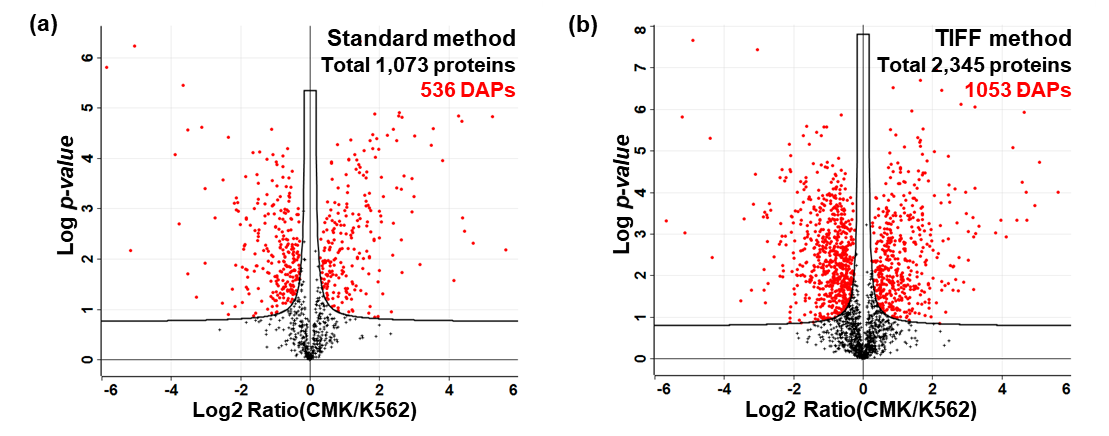

**Figure S6.** **(a-b)** Statistics analysis to identify differentially abundant proteins (DAPs) between CMK and K562 cells using iBAQ intensities (t-test FDR < 0.05 and S_0_ = 0.1). Volcano plots for **(a)** standard method and **(b)** the 4-CV TIFF method. Total quantified proteins and DAPs were labeled with red color.

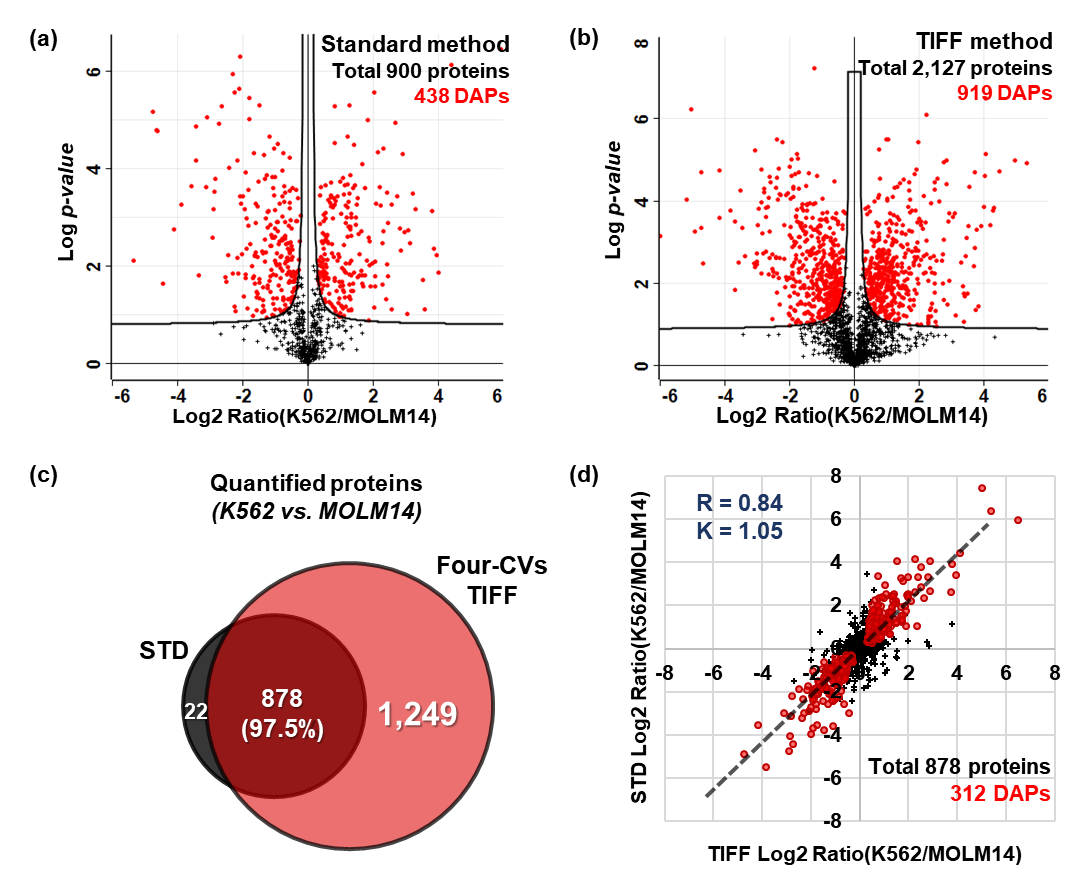

**Figure S7.** (**a-b)** Statistics analysis to identify differentially abundant proteins (DAPs) between K562 and MOLM14 cells (t-test FDR < 0.05 and S_0_ = 0.1). Volcano plots for **(a)** the standard and **(b)** 4-CV TIFF methods. **(c)** Overlap of quantifiable proteins between K562 and MOLM14 cells measured by standard and TIFF methods (4 CVs). **(d)** The linear correlation and slope of log2 transformed fold changes of K562 and MOLM14 proteins between the 4-CV TIFF and STD methods. Red dots indicate DAPs in both methods calculated by t-test (FDR<0.05, S_0_=0.1).

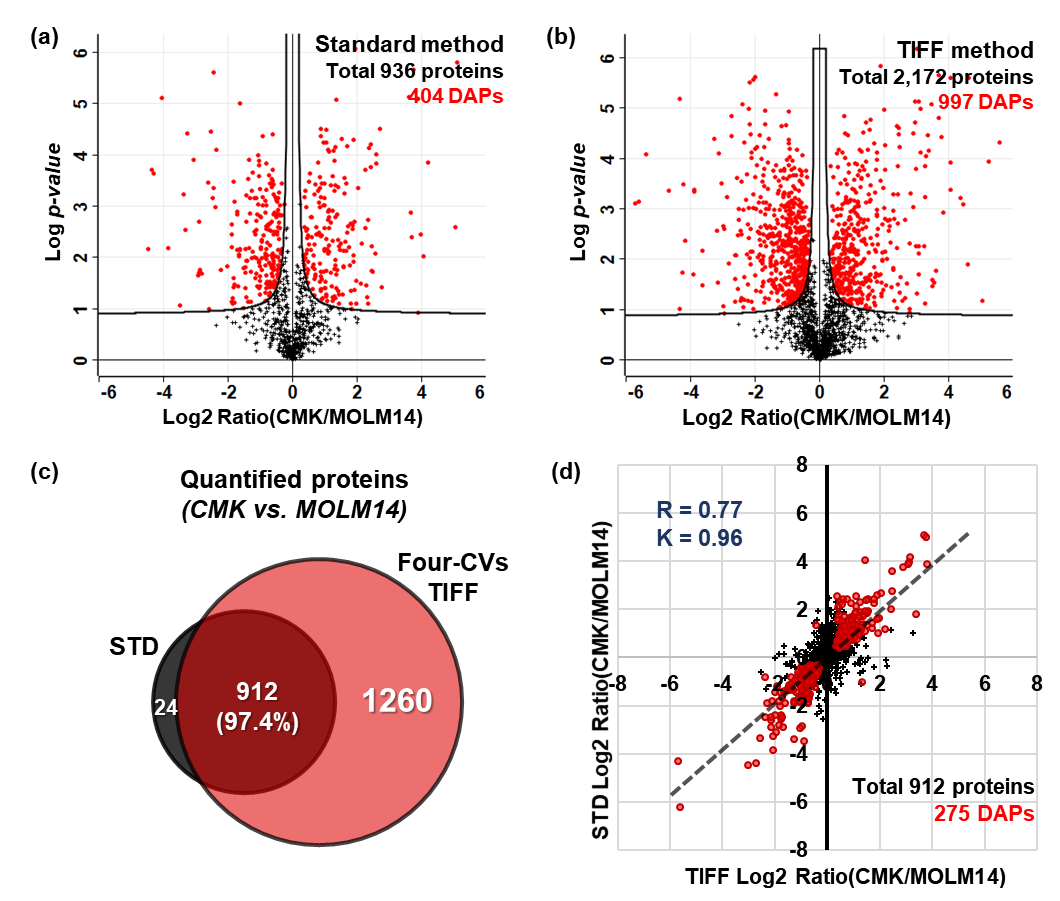

**Figure S8.** **(a-b)** Statistics analysis to identify differentially abundant proteins (DAPs) between CMK and MOLM14 cells (t-test FDR < 0.05 and S_0_ = 0.1). Volcano plots for **(a)** the standard and **(b)** 4-CV TIFF methods. **(c)** Overlap of quantifiable proteins between CMK and MOLM14 cells measured by the standard and 4-CV TIFF methods. **(d)** The linear correlation of log2-transformed fold changes of CMK and MOLM14 proteins between the 4-CV TIFF and STD methods. Red dots indicate (DAPs) calculated by t-test (FDR<0.05, S_0_=0.1).

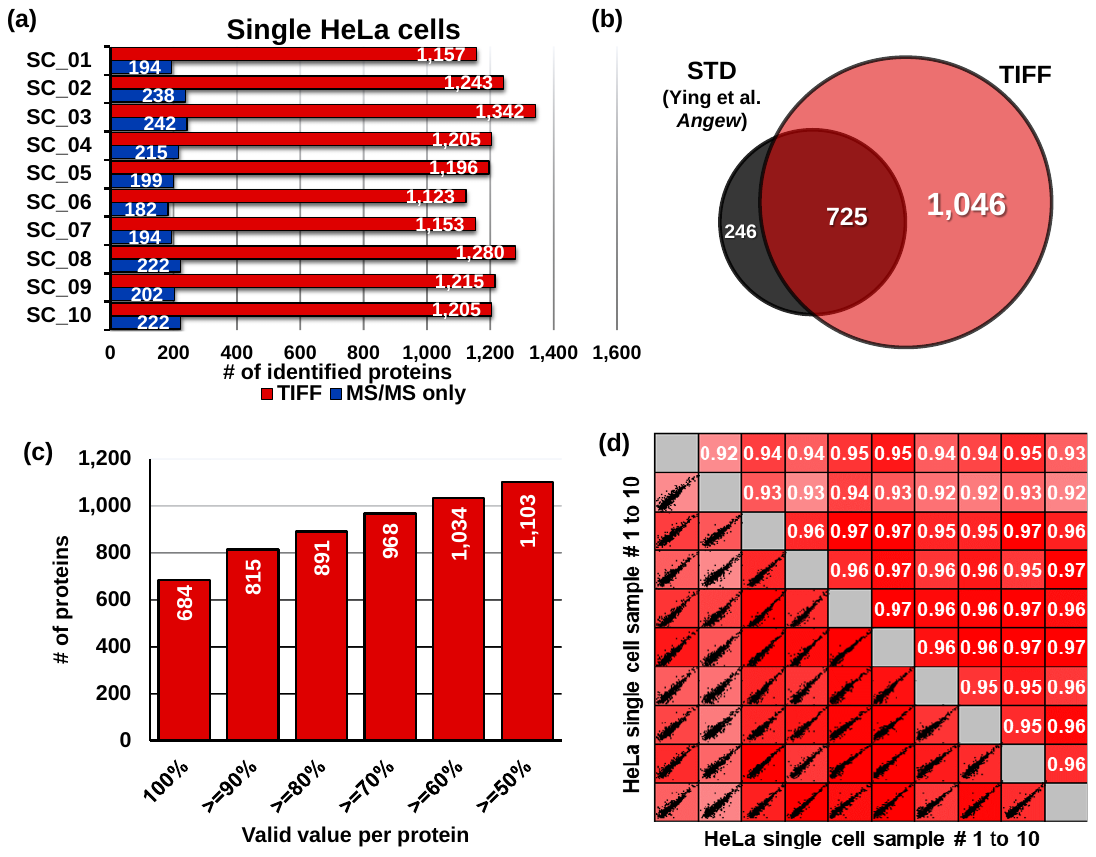

**Figure S9. ScProteomics of HeLa cells using TIFF method. (a)** Number of protein groups in single HeLa cells identified by MS/MS only (blue) and by the 4-CV TIFF method (red). **(b)** The overlap of identified proteins in single HeLa cells obtained in this study and a previous study with a similar LC-MS setting but without FAIMS ^2^. **(c)** The numbers of proteins having valid values from 50% to 100% across the 10 single cells. **(d)** Pair-wise correlations of protein iBAQ intensities between the 10 cells. Proteins containing >70% valid values were required.

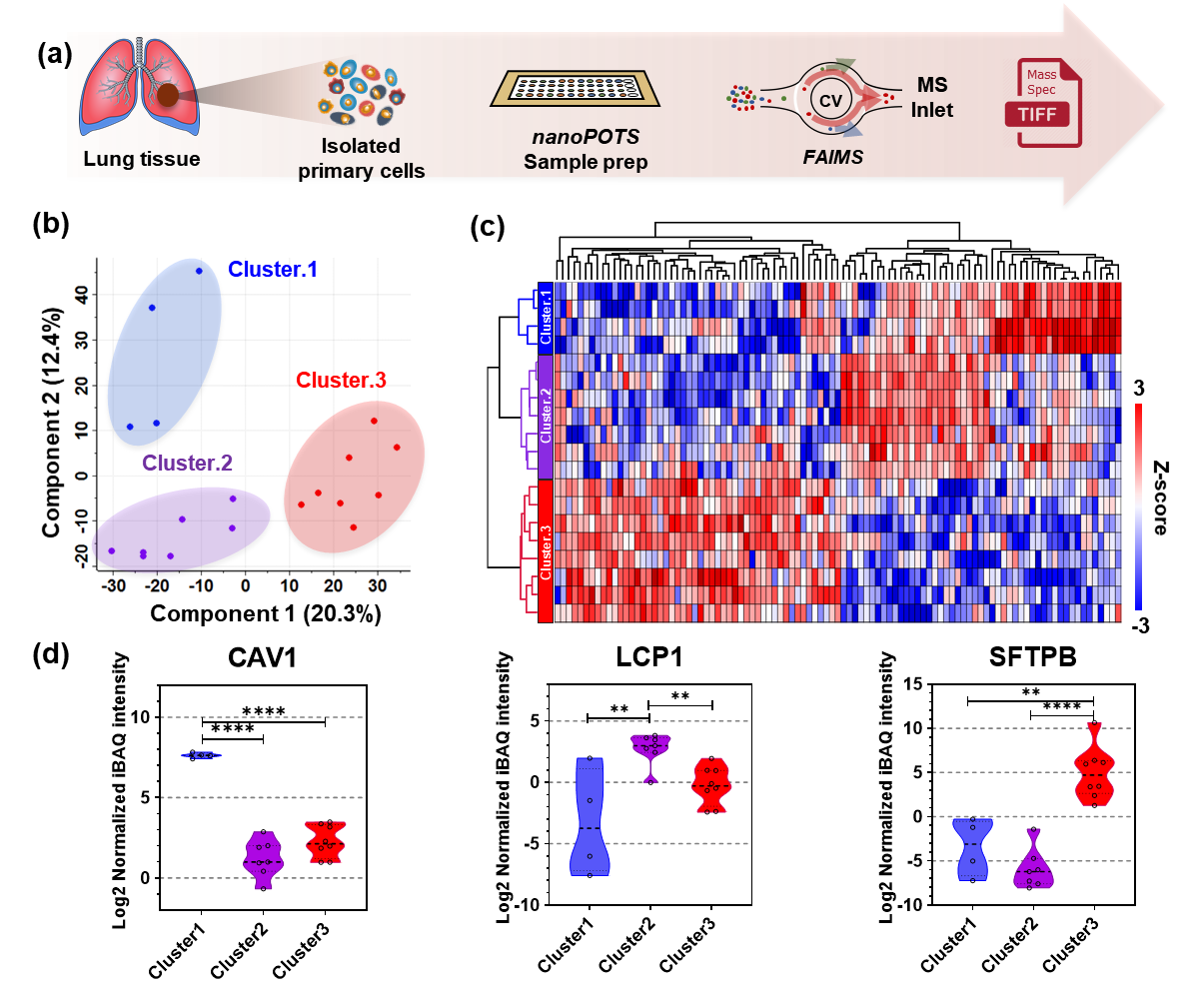

**Figure S10. ScProteomics for classifying cell populations of a human lung.**

**(a)** Schematic workflow for scProteomic analysis of dissociated human lung tissue from a 2-year old donor. The tissue was dissociated into the single cells followed by applying the scProteomics pipeline including FACS isolation, nanoPOTS processing, autosampler-LC, and TIFF MS method. **(b)** PCA plot of un-defined cell types. **(c)** Heatmap of differentially abundant proteins (DAPs) by ANOVA test. **(d)** Representative proteins of putative cell types. Cluster 1, 2, and 3 were predicted as lung endothelial, immune, and epithelial cells, respectively (**<0.01, ****<0.0001).

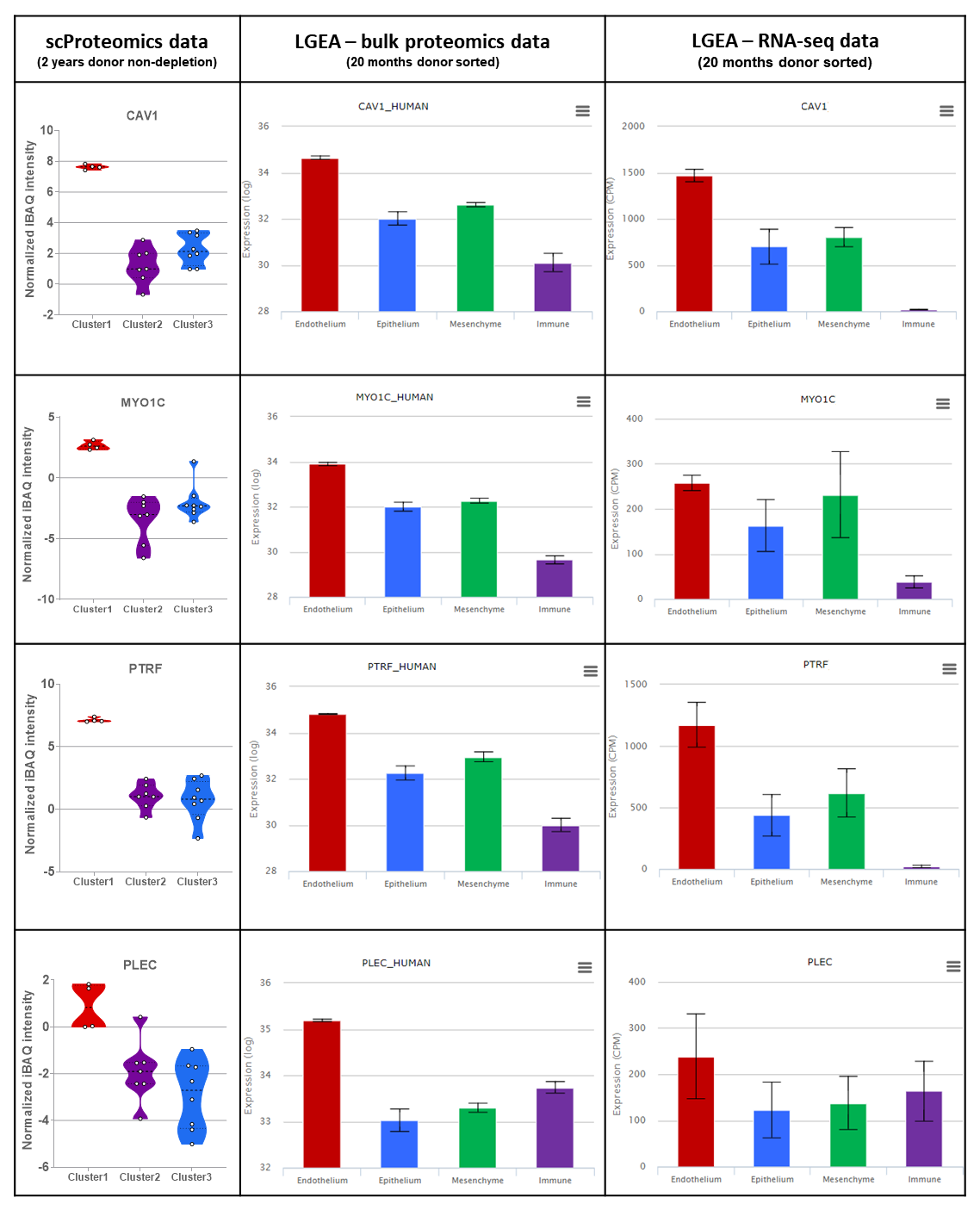

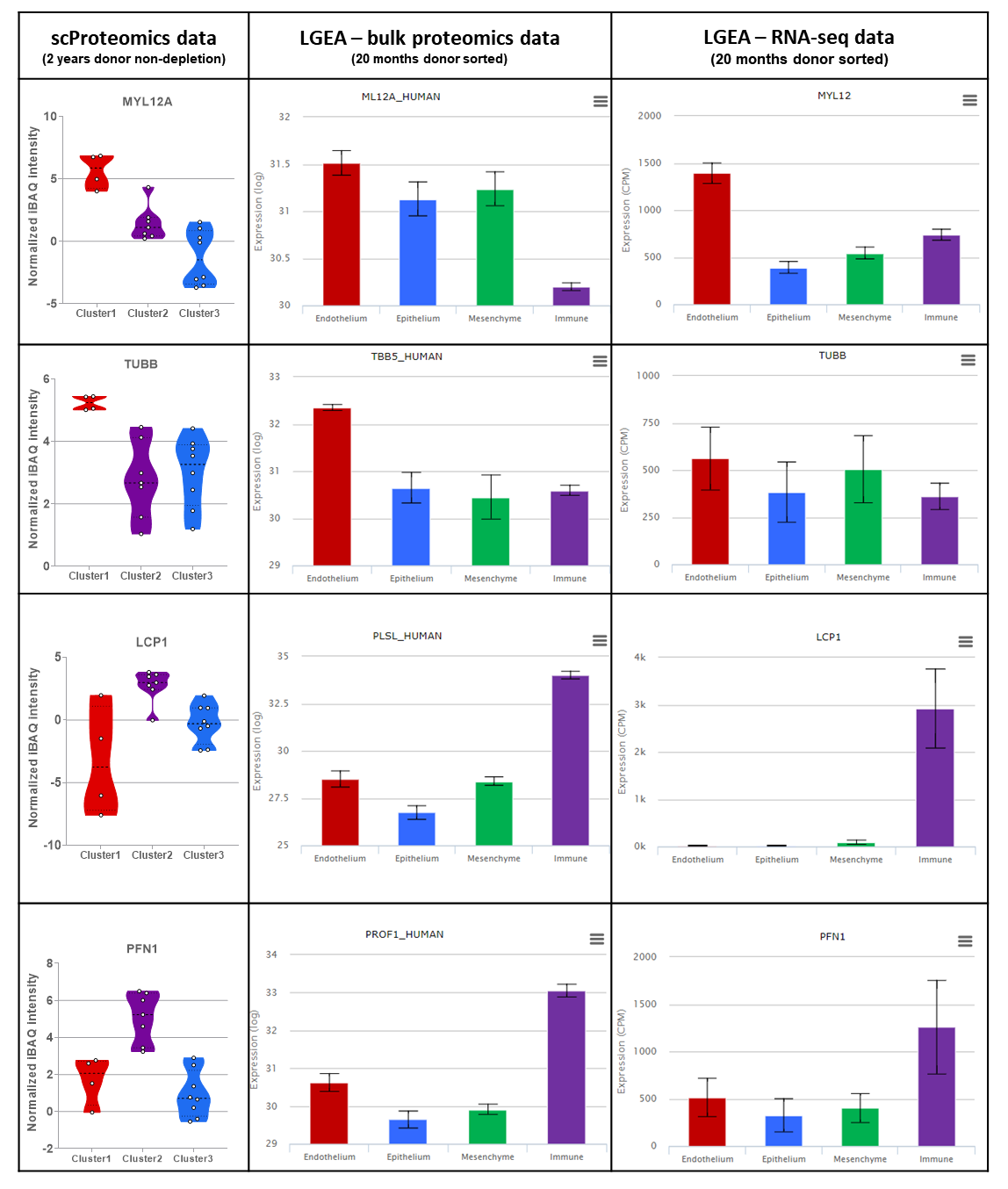

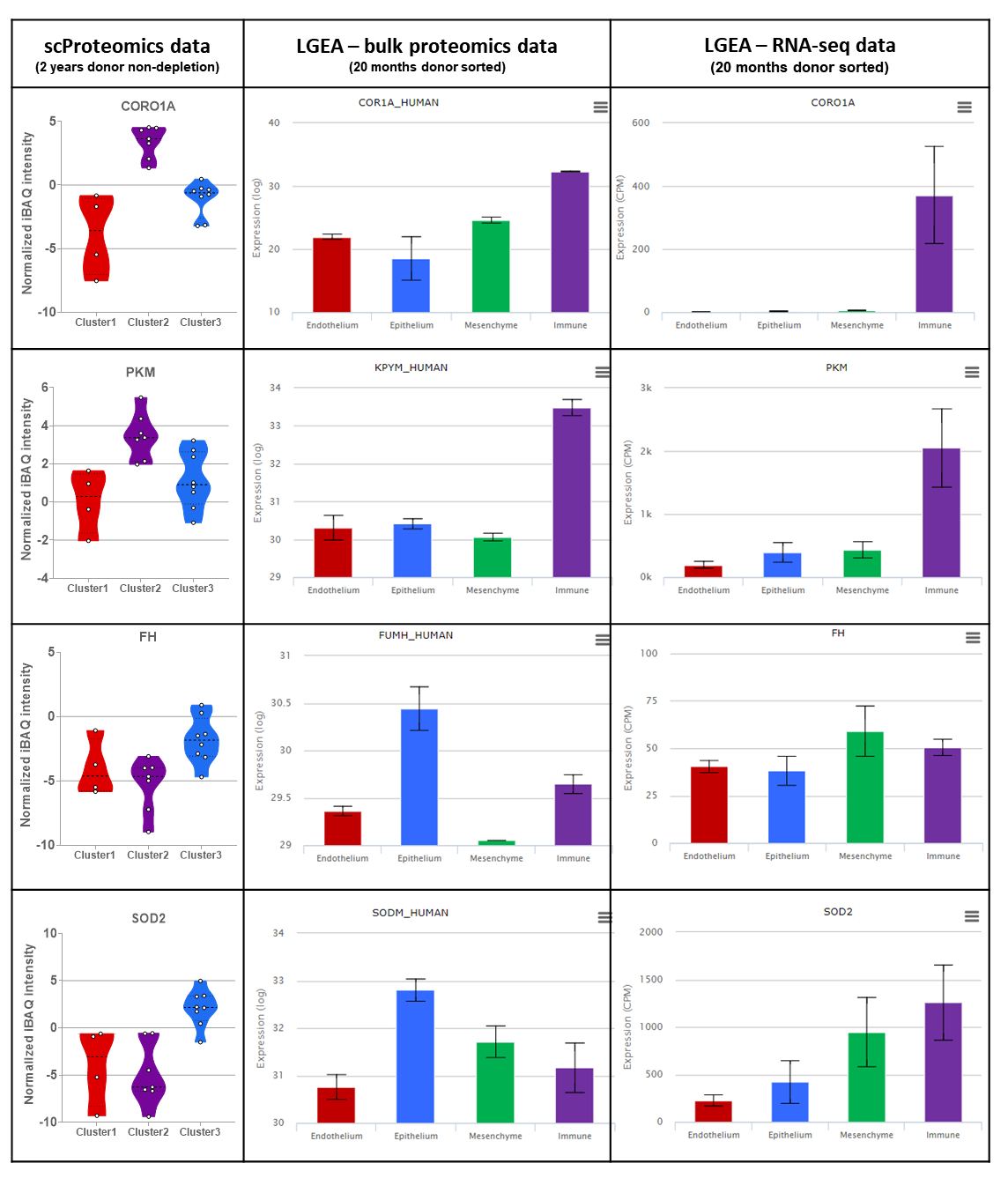

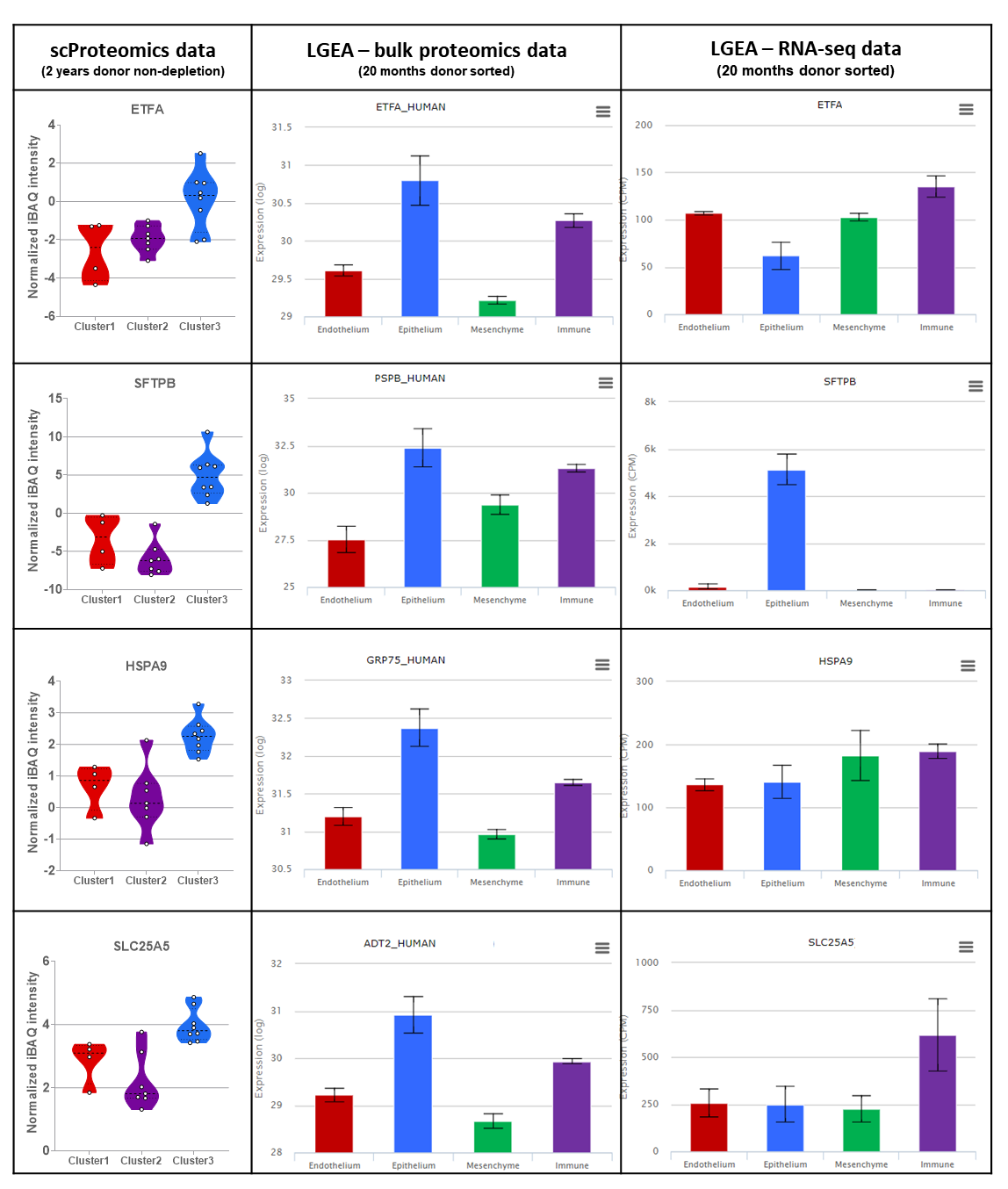

**Figure S11.** The abundance distributions of representative proteins markers in the scProteomics data and lung gene expression analysis (LGEA) database (<https://research.cchmc.org/pbge/lunggens/mainportal.html>) containing sorted human lung endothelial, epithelial, immune and mesenchymal cells measured by bulk proteomics and RNA sequencing ^3^.

| **Single-cell approach** | **LC system**  **(column I.D. in µm, flow rate in nL/min)** | **MS instrument** | **Cell Type** | **Protein Groups  (by MS/MS)** | **Proteins Groups (MBR)** | **Reference** |
| --- | --- | --- | --- | --- | --- | --- |
| nanoPOTS | 30 / 50 | Lumos | HeLa | 211 | 669 | Zhu et. al, *Angew chem*, 2018 ^2^ |
| nanoPOTS with autosampler | 50 / 150 | Lumos | MCF10 | 250 | 773 | Sarah et. al. *Analytical Chemistry*, 2020 ^4^ |
| nanoPOTS with narrow-bore LC | 20 / 20 | Eclipse | HeLa | 362 | 874 | Cong et. al. *Analytical Chemistry*, 2020 ^5^ |
| nanoPOTS with FAIMS | 20 / 20 | Eclipse | HeLa | 683 (By MQ)/  1056 (By PD) | 1475 | Cong et. Al. *Chemical Science*, 2021^1^ |
| nanoPOTS with TIFF | 50 / 100 | Lumos | HeLa | 209 | 1212 | *This study* |

**Supplementary Table 1**. Numbers of identified proteins in single mammalian cells from previously published papers using nanoPOTS and label-free analysis.

Proteins were identified by MS/MS or by matching between runs (MBR) algorithm in MaxQuant software ^6^ (MQ: MaxQuant, PD: Proteome Discoverer).

**Supplementary references**
